# Improved Modeling of Droplet Motion in Open-Format Digital Microfluidic Devices

**DOI:** 10.1101/2022.11.30.518563

**Authors:** Karanpartap Singh, Benjamin G. Hawkins

## Abstract

Electrowetting is an electrokinetic effect whereby an applied electric field induces changes in the measured contact angle at a fluid-surface contact line. On hydrophobic, dielectric electrode surfaces, this effect generates droplet motion termed “electrowetting on dielectric” or EWOD. Applications of this phenomenon range from lab-on-a-chip to liquid lenses capable of altering their topology and focus within milliseconds. Electrowetting or EWOD theoretical models quantifying this effect fall into two paradigms: the Young-Lippman and the electromechanical theories. In this work, both paradigms were simulated to predict the velocity of a water droplet moving over an array of electrodes. Results were compared to experimental observations of measured velocities for two dielectric films: ETFE and household cling film. Theoretical model parameters, namely the length scale of the Maxwell force on the droplet, were also determined to align simulation and experiment. The results reveal the trend of droplet velocity in relation to applied voltage, and recapitulate the relationship between the two models.

## 1 INTRODUCTION

Electrowetting refers to the electric-field-induced change in contact angle between a solid planar substrate and a fluid droplet. Typically, an aqueous droplet and a hydrophobic substrate are used in either a hydrophobic or gaseous third phase. This configuration – often referred to as “electrowetting on dielectric,” or EWOD – enables a range of fluid manipulation strategies collectively termed “Digital Microfluidics” (DMF). EWOD effects can be used to electrically transport micro- or nanoliter-sized fluids with high precision and speed, minimal human contact, and low power consumption, with numerous applications [1]. These benefits over other droplet actuation methods such as magnetic [2] or acoustic [3] make electrowetting an important tool in microfluidics, variable-focus lenses, and other MEMS devices [4, 5].

The two leading models describing DMF droplet actuation are thermodynamic and electromechanical in nature, and produce the same results for the balance between the interfacial forces of the droplet [4], though with differing implications. While the thermodynamic model quantifies the contact angle of the droplet and its deformation at high voltages, the electromechanical model describes only the force on the droplet. Additionally, given that this force is independent of the shape of the droplet, the EM model suggests that the deformation and movement of the droplet can be considered separately [6].

### 1.1 Conventional electrowetting model

The thermodynamic model of electrowetting balances the surface tensions at liquid, solid, and vapor interfaces between a wetting droplet and a surface. In 1875, Lippmann observed that capillary forces can be controlled with external electrostatic fields [7]. He noted that the effective change in surface tension is proportional to the applied potential squared, represented in Equation 1.

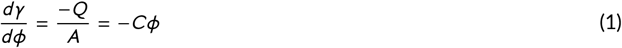

Where *γ* is the surface tension at the liquid-solid interface, *Q* is the induced surface charge, *A* is the interfacial area, and *ϕ* is the local electric potential. The ratio *Q* /*A* is defined as a capacitance, *C*. Assuming a known surface tension in the absence of an applied electric potential, *γ*_0_, the equation reduces to Equation 2:

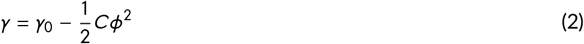

Meanwhile, Young showed that the contact angle of a droplet depends on the surface tensions at its triple-contact point [4]. This relationship is shown in equation 3 as a function of the constituent surface tensions between the solid (S), gas (G), and liquid (L).

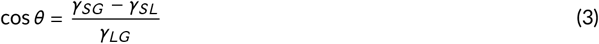

In the 1990s and early 2000s, Berge combined these relationships to produce the Young-Lippman model, which has become convention for electrowetting theory [8]. Treating the dielectric layer as an ideal parallel-plate capacitor, equation 4 is obtained to describe the contact angle of a droplet based on its initial contact angle (*θ*_0_), dielectric constant (*ϵ*_*r*_), applied voltage (*V*), liquid-vapor surface tension (*γ*), and dielectric layer thickness (*d*_*f*_).

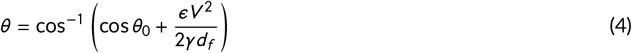

The motion of a liquid droplet in response to the above phenomenological description of electrically actuated change in the interfacial contact angle arises from a propagation of stresses across the surface of the droplet originating from local and non-uniform changes to the contact angle. The droplet moves in response to non-uniform surface stresses until a new thermodynamic equilibrium point is reached [9].

### 1.2 Electromechanical model

Another model for electrowetting, first proposed by Jones, examines the electromagnetic field and force resulting from the applied voltage [10]. We obtain the net force by integrating the flux density of the electric fields over the surface of the droplet via the Maxwell stress tensor. The time-averaged Maxwell stress tensor, 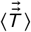 assuming a steadystate AC electric field is given in Equation 5 and the resulting time-averaged electromechanical force acting on the droplet, 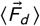, ignoring osmotic contributions, is given in Equation 6.

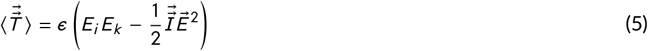

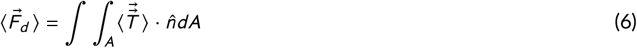

where 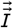 is the second-order isotropic tensor and 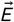 is the electric field. Following the work of [11], the net electromechanical force in the x-direction acting on the outer boundary of the droplet after performing this integration, tangential to the electrode surface, is found in Equation 7. Note that the Maxwell stress is concentrated in a small region on the edge of the droplet, and decays rapidly away from the triple contact line owing to the dependence on *V* ^2^.

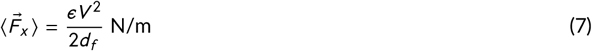

where 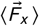is the time-averaged force in the x-direction on the droplet, *V* is the potential difference applied between neighboring electrode pads, and *d*_*f*_ is a characteristic decay length of the field and resultant force normal to the electrode along the triple contact point.

### 1.3 Recent developments

Though the two discussed models define the contact angle change and forces on the droplet for reasonable voltages, phenomenon beyond a certain critical voltage are not yet fully understood. Experiments and simulations show clear saturation of the contact angle when exceeding this critical voltage, though the physical mechanism of this saturation is inconclusive [12]. Additionally, the contact angle is experimentally found to converge to the same value at high voltages regardless of factors such as dielectric thickness or type, further complicating this problem and opening the door for more conclusive theories [13], [14]. Recent work has also shown an effect of velocity saturation in two-plate microfluidic devices [15], shedding more light on the nature of the resistive forces that oppose droplet motion. We do not observe this effect in our open, one-plate device, likely due to the low values of our applied voltages.

On the application front, recent developments in superhydrophobic surfaces allow much larger contact angle changes in electrowetting devices. The use of AC electric fields can also allow oscillation of the droplet or even self-propulsion [16]. Combining superhydrophobic surfaces with AC electric fields leads to behaviors such as droplet bouncing [17], providing a pilot for droplet transportation with anti-gravity-like effects. Advancements in fabrication techniques have also made electrowetting more accessible than ever, with inkjet printing techniques capable of producing functional and disposable devices for as low as $0.63 per device, making clinical applications feasible [17].

In the present study we conducted finite element simulations of electrowetting phenomenon to compare the thermodynamic and electromechanical models. In the thermodynamic model, we considered only the deformation of the droplet due to the contact angle change at its triple-contact point. In the other, we assumed the shape-independence of the electromechanical force on the droplet following [6] and did not explicitly model deformation or the change in the droplet’s contact angle. In both cases, we tested and quantified the parameters required to balance the results of these models. Specifically, we confirm that the Maxwell stress on the droplet is localized to the triple point on the contact line on a length scale about 10 times larger than the dielectric thickness. We show a strong correlation between the droplet velocities obtained from these two models as well as with experimental results obtained on an open-configuration digital microfluidics platform.

## 2 METHODS

We modeled single droplet, open format electrowetting in two dimensions using COMSOL Multiphysics. Separate simulations were created to test the Young-Lippman and electromechanical theories, and the obtained results were compared to experimental measurements taken on the open-source OpenDrop digital microfluidics platform [18]. Finally, parameters such as Navier slip length and electromagnetic force decay length required to match the results of the two models were quantified.

### 2.1 Simulation setup

#### Geometry

The simulation geometry, shown in Figure 1, is modeled after the OpenDrop device to match our experimental setup: four gold-coated 2.75mm electrodes separated by 0.1016 mm (4 mils) are simulated in an open configuration and surrounded by air. The 2D simulation includes an out-of-plane thickness equal to the width of the square electrodes (2.75 mm). A 10 μL spherical droplet of water with a radius of 1.6839 mm is placed on the center of the starting electrode and is given an out-of-plane thickness equal to the diameter of the droplet. It rests on top of a 12 *μ*m thick ethylene-tetra-fluoro-ethylene (ETFE) film, which extends over the electrodes. Another dielectric film, created from household cling film and coated with a hydrophobic layer of peanut oil, was also tested. This film was modeled as low-density polyethylene (LDPE). The properties of both of these films are included in the supplementary data.

**FIGURE 1.**
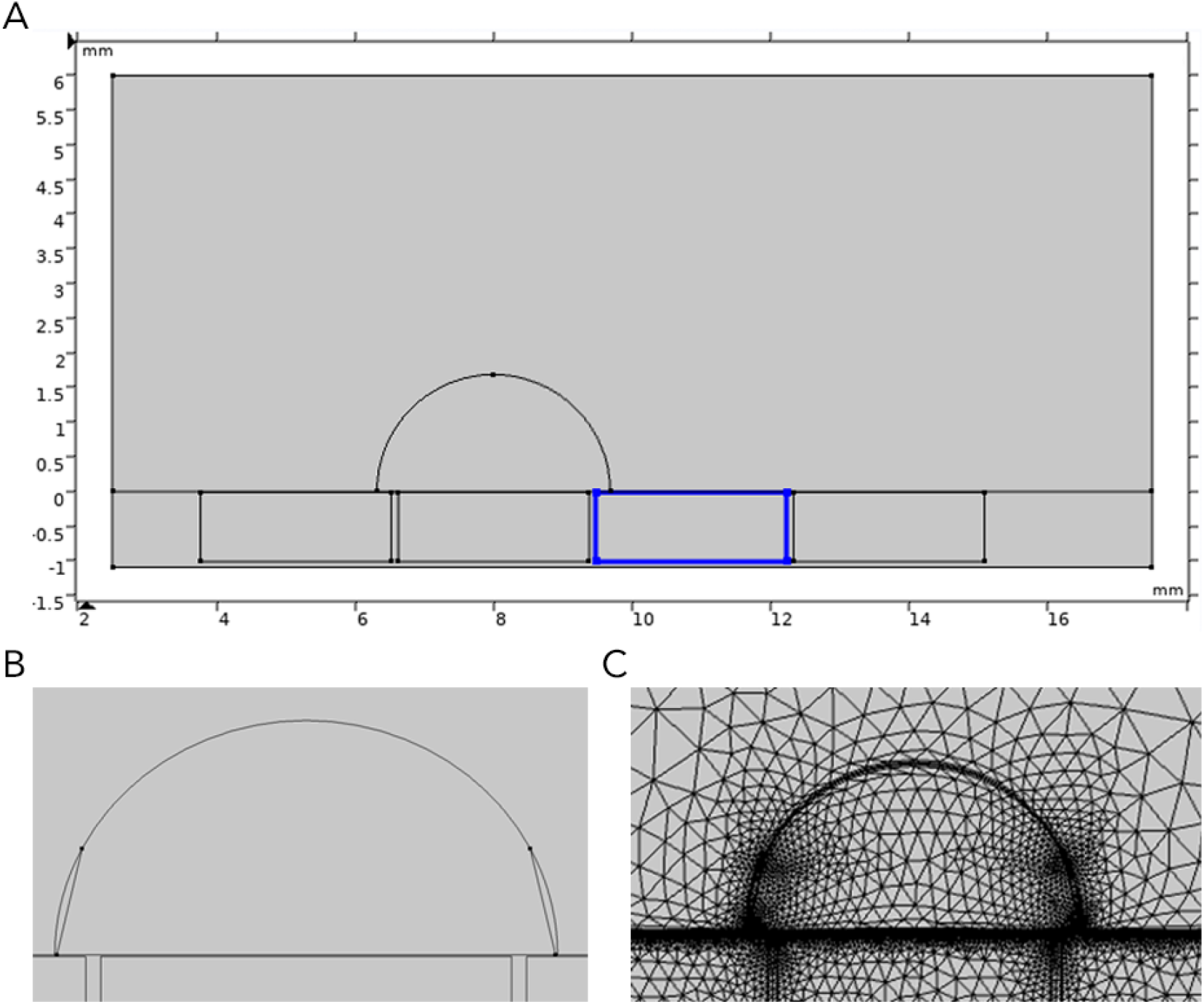
A) Simulation geometry. B) Droplet partitioning for electromechanical simulation model. C) Simulation mesh.

The fluid and solid domains are built separately and joined as an assembly, with continuity boundary conditions enforced between them in fluid and electrical physics modules. Separating the domains was necessary to accommodate the moving mesh and mesh slip conditions at the interface.

#### Fluid Physics

The interactions between the droplet and the air surrounding it are modeled using a Creeping Flow and Electric Currents nodes in COMSOL Multiphysics. The flow is modeled as weakly compressible with gravity included and inertial terms ignored. The boundary between the droplet and dielectric is given a Navier slip wall condition with a slip length of 43 nm, consistent with atomic force microscopy measurements on hydrophobic films [19]. The impact of this parameter was also tested by varying it between 30 and 70 nm and observing the resulting droplet velocities. Remaining fluid boundaries were modeled as open boundaries with a normal stress of 0 N/m^2^.

Since the initial contact angle and shape of the droplet are not known, the initial geometry only approximates the initial equilibrium shape of the droplet. To determine the equilibrium shape of the droplet, prior to electrical actuation, the gravitational force was applied to the droplet in the absence of other effects and the moving mesh allowed to reach equilibrium. This state was then used as the initial condition for electrical actuation. COMSOL’s Events interface was used to apply the gravitational force at 0.2*s* and the electrical force at 0.5*s*.

#### Electrical Physics

COMSOL’s Electric Currents node solved for the electrical potential in all domains of the simulation. The actuation voltage is applied to the third electrode, and the other electrodes are kept grounded. Actuation voltages were varied using a parametric solver. Other boundaries were assigned an Electric Insulation boundary condition (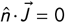, where 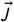 is the electric current). Electrical properties, namely conductivity and permittivity, were taken from COMSOL’s built-in material library, measured, or sourced from literature and are included in supplementary data.

#### Thermodynamic Model

For the thermodynamic model, the droplet-air interface was modeled using a Fluid-Fluid Interface node (water/air), with a contact angle specified by Equation 4 at the droplet-solid-air interface. The contact angle depends on the local electric potential determined by an Electric Currents node. In response to changes in the contact angle, the droplet must deform. Surface tension along the droplet-air interface transfers deformation along the interface and drives droplet motion until a new equilibrium state is found centered on the actuated electrode.

#### Electromechanical Model

For the electromechanical model, the simulation is kept primarily the same with the contact angle now set to the film’s known contact angle with water. A volume force is applied to a small section at the edge of the droplet with the net force magnitude in equation 7. The size of this slice was adjusted to match the results of the Young-Lippman model and therefore determine the physical length-scale of the force on the droplet. The droplet is partitioned on both sides to avoid meshing inconsistencies (see Figure 1). However, the volume force is only applied to the side adjacent to the activated electrode.

#### Mesh

Meshing is done with different resolutions for the components - the dielectric film, electrodes, and droplet - of the simulation, and tailored to balance simulation time and accuracy. Finer mesh presets were chosen for the droplet and dielectric film, as shown in Figure 1c, while coarser meshes could be used for the electrodes and areas of air far away from the droplet. The simulated droplet velocity was lower with coarser mesh presets, and stabilized to the reported values with the ’Extra Fine’ or ’Extremely Fine’ options. A boundary layer mesh also surrounds the droplet to ensure adequate accuracy in the crucial water/air boundary.

The moving mesh module was used to capture droplet deformation and movement. A Deforming Mesh node was applied to the fluid domains (air and droplet). Yeoh smoothing and a stiffening factor of 10 allow deformation of the air and droplet domains. The fluid and electrode/substrate domains are joined by a Mesh Slip boundary condition.

### 2.2 Device fabrication and use

We chose the OpenDrop device for its affordability, ease of fabrication, and voltage range of 150-300 V. Parts to fabricate the device were sourced from PCBWay [20] and Mouser Electronics [21]. A crucial consideration was the tolerance between the electrodes, which is 4 mils or 100 μm on the device, and the appropriate PCB manufacturing service had to be chosen for successful fabrication. The OpenDrop is prepared for use by first cleaning the surface of the electrodes with 70% isopropanol and allowing them to air dry.

After an application of 10 μL of mineral or peanut oil, the dielectric film is carefully placed on top and pressed down to remove air bubbles. This film is coated with a hydrophobic layer to complete the setup. These steps are demonstrated for the LDPE dielectric film in Figure 2. Variations in these processes could be reduced through precise applications of the hydrophobic layer and films in a clean-room, though this was not done in this study due to limited access to these facilities.

**FIGURE 2.**
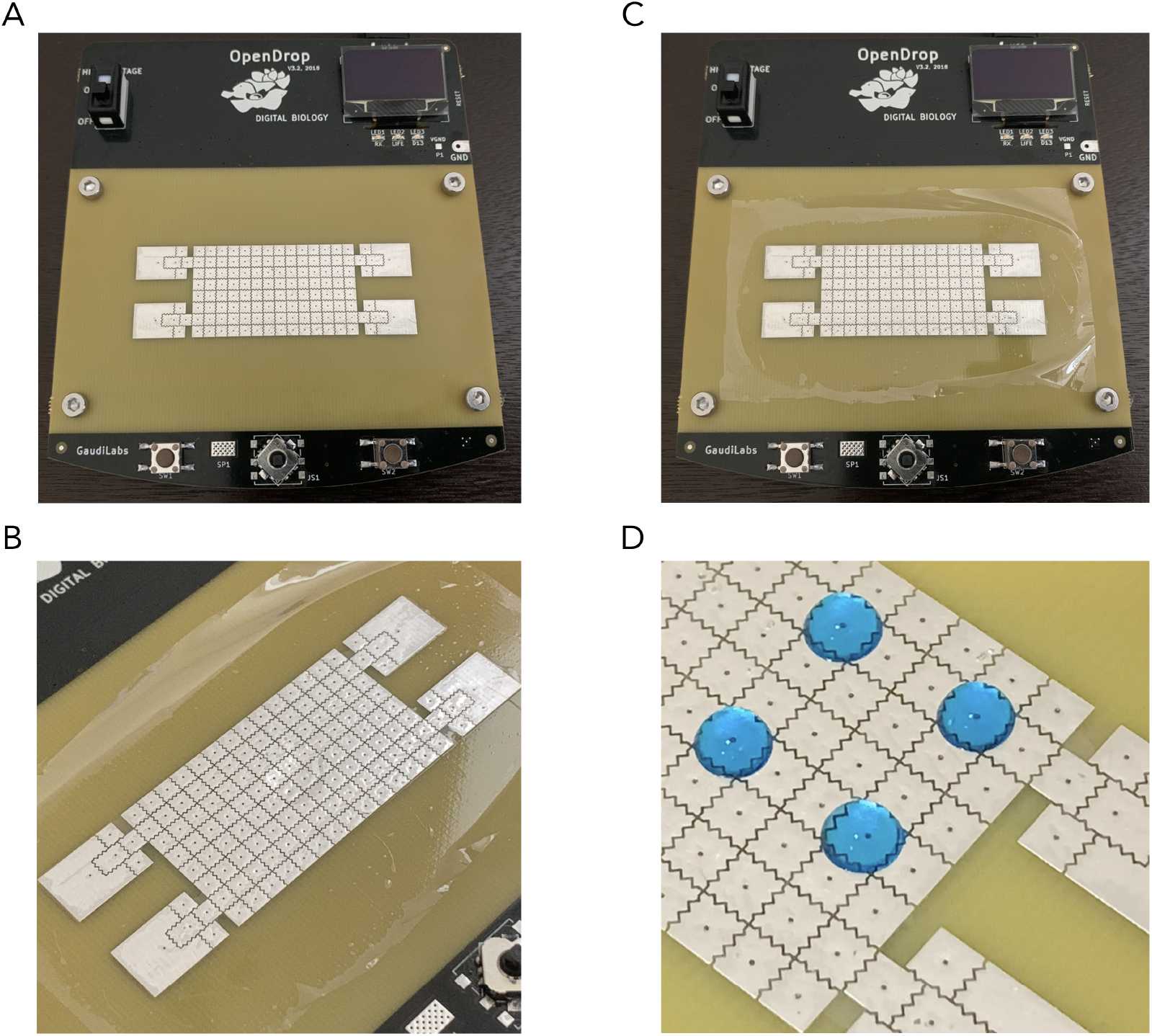
Device preparation for LDPE and peanut oil film. A) Cleaned device with layer of oil. B) Applied dielectric film (cling film). C) Layer of peanut oil on top of film. D) Prepared device with droplets.

The fluid used in the velocity measurements is a simple combination of ink and distilled water, of which 10 μL is pipetted onto the device to create each droplet. After placement on the device, a small piece of wire is connected between the ground terminal on the OpenDrop and the droplet to remove any accumulated static charge.

A proper hydrophobic coating on top of the dielectric film was observed to be a crucial component to achieving smooth droplet motion. The ETFE films used in this study were manufacturer-coated and sourced from AGC Chemicals [22]. The contact angle for the ETFE films was measured (VCA Optima) as 105.5° (*σ* = 1.96°). Though not tested in this work, an additional hydrophobic coating such as FluoroPel may be used [23]. When using household cling film, peanut oil was used, and produced a large contact angle and consistent droplet motion. The contact angle of the native household cling film was measured (VCA Optima) as 103.2° (*σ* = 6.1°).

### 2.3 Velocity measurement

Droplet velocity measurements were made by filming the motion at 1080p and 240fps using an iPhone XS (see Figure 3 in Results). The time taken to travel across one full electrode was observed and used, along with the length of the electrode, to compute the droplet’s velocity. The droplet velocity was averaged between two trials for each film and applied voltage. Similarly, for the simulations, the time for the right edge of the droplet to move the length of one electrode was measured.

**FIGURE 3.**
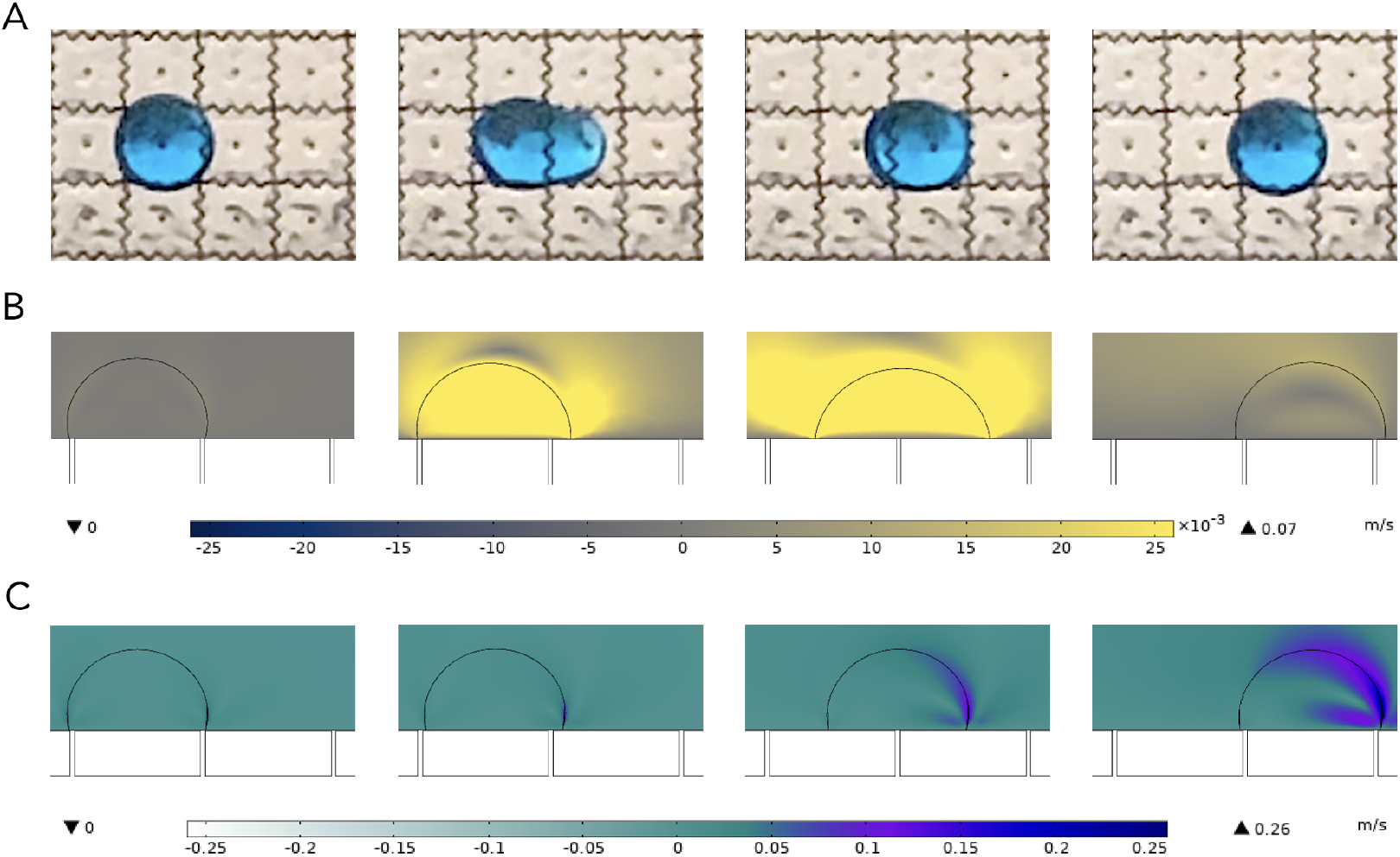
A) Initial deformation and subsequent motion of the droplet in an open electrowetting configuration. B) Young-Lippman model simulation with only deformation and contact angle change modeled. Colors and color bars represent the velocity fields of the droplets. C) Electromechanical model simulation with only the electromechanical force on the droplet modeled.

## 3 RESULTS

## 4 DISCUSSION

Figure 3b displays the simulated deformation of the droplet under the Young-Lippman model for the ETFE film and is consistent with the experimental observations in Figure 3a. Upon activation of the electric field, the local contact angle changes according to the Young-Lippman model. The initial deformation of the droplet towards the actuated electrode propogates surface stresses across the droplet and propels the fluid forward until it reaches the next grounded electrode. At this point, the contact angle settles to its resting value, and motion ceases. No external forces (other than gravity) were simulated here, and the change in the contact angle between the droplet and planar surface was the sole driver of droplet motion. In the electromechanical model, there was little to no deformation of the droplet, as seen in Figure 3c. The calculated total force from Equation 7 was applied to a segment of the droplet-air interface originating at the triple point.

Referring to Figure 4, the simulated and experimental droplet velocities agree well with each other. For the ETFE dielectric film, a maximum droplet velocity of 28.6 mm/s was observed at 300V for the experimental trials and both simulation models. Across the voltage range, the simulated droplet velocities remain within an error of 20% from our experimental measurements. More variation is observed with the LDPE dielectric film.

**FIGURE 4.**
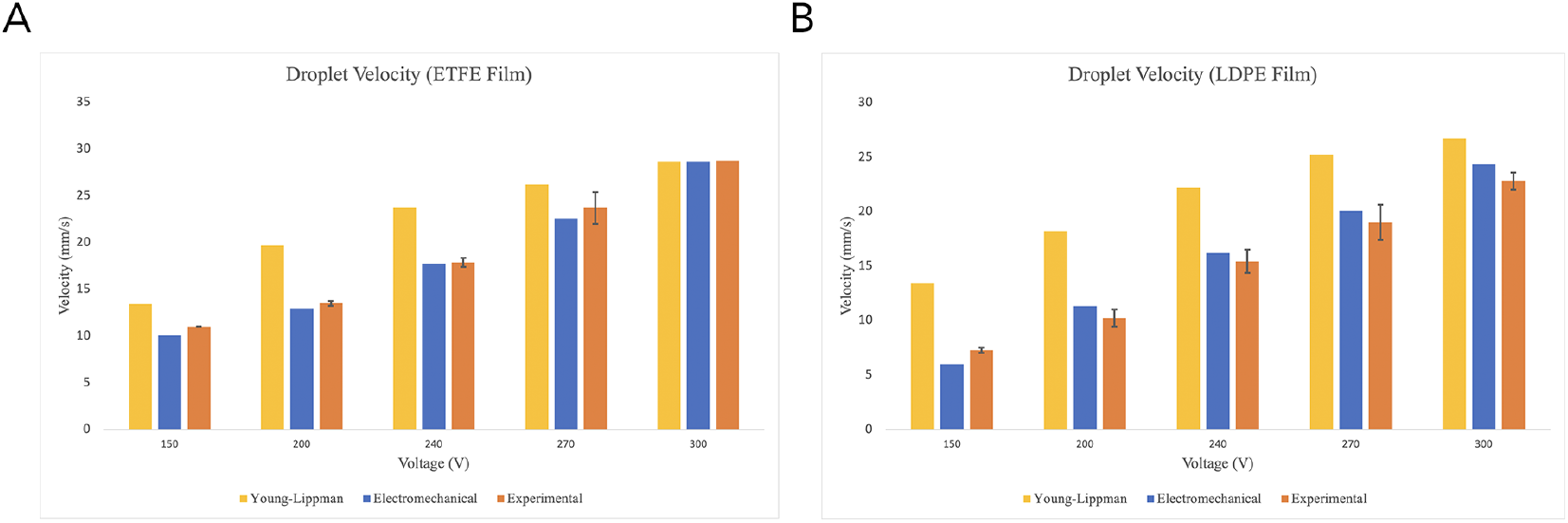
A) Experimental (n = 2, mean ± s.e.m) and modeled droplet velocities for an ETFE dielectric film. B) Experimental and modeled droplet velocities for an LDPE dielectric film.

In describing the electromechanical model, the Maxwell stress on the droplet is assumed to decay to a negligible value beyond a few *d*, the dielectric thickness, from the triple-contact-line [11, 4]. To quantify the exact length scale of this decay, we adjusted the size of the domain on the edge of the droplet to which the stress was applied until droplet velocities agreed with the thermodynamic model. This was determined to be on the order of 10*d*, which is consistent with the approximate measure given in previous studies.

Experimentally, the combination of household cling wrap and peanut oil performed surprisingly well for its accessibility and affordability, yielding good durability across trials and consistent motion at speeds of up to 22.7 mm/s at 300 V. Simulation results for this system indicate a maximum droplet velocity of 24.3 mm/s for the electromechanical model and 26.7 mm/s for the thermodynamic. Some of the observed variations between the models and experimental results could be a result of the method used to measure the droplet velocity. At the utilized frame rate of 240fps, a difference of a single frame leads to an error of 2-3 mm/s in the resulting velocity, or 8-12%.

Prior work has quantified the velocity of a smaller 2.5 μL droplet on a 13μm ETFE film in an open configuration. Despite the similarity in experimental setup, this study reports a maximum observed droplet velocity of 6.0 mm/s at 295 V [24]. This difference is likely a result of the different methodology used to make these measurements, as well as the smaller volume of the droplet. Our velocity results are an average across the entire length of one electrode, while previous work takes the average velocity over the initial 100 μm gap between the electrodes. The smaller velocities are thus in-line with theoretical results. As the droplet covers more of the actuated electrode, the force acting on it increases, making it speed up. Similarly, if there is no part of the droplet’s contact line on the electrode, there is no force or motion [25].

Figure 5 displays the dependence of the simulated droplet’s velocity on the chosen Navier slip length for the thermodynamic model, as well as the decay length of the force on the droplet for the electromechanical model. While we use the slip length measured in prior literature [19], this parameter can be varied to achieve simulations closer to experimental observations. In the electromechanical model, the decay length scale of the Maxwell force on the droplet can be similarly varied to scale the droplet’s velocity.

**FIGURE 5.**
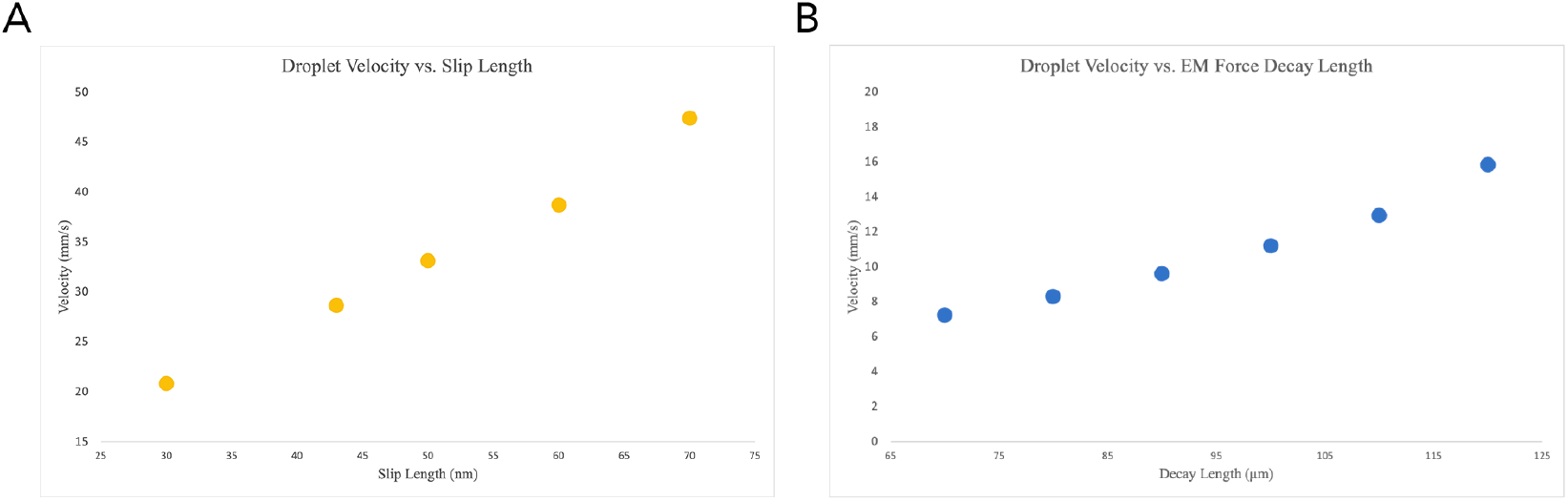
A) Droplet velocity predicted by thermodynamic simulation vs. Navier slip length between droplet and film surface for an ETFE film at 300V. B) Droplet velocity predicted by electromechanical simulation vs. decay length-scale of EM force for an ETFE film at 200V.

## 5 CONCLUSION

In this paper, we developed simulations of the electrowetting effect, considering separately motion described by the Young-Lippman and the electromechanical model. We showed reasonable agreement between experimental results and the droplet velocities predicted by both models, and described the trend of these velocities in open-format electrowetting. Additionally, we quantified the length scale of the Maxwell force on the droplet to be on the order of 10 times the dielectric thickness. The Navier slip length required between the droplet and surface for accurate simulation was also verified to be 43 μm. These findings shed light on the relationship between these two commonly applied models for open electrowetting, and could allow better modeling of this effect.

## Abbreviations

DEP: Dielectrophoresis
EWOD: Electro-Wetting on Dielectric
DMF: Digital Micro-Fluidics
MST: Maxwell Stress Tensor.

## Supporting Information

**TABLE 1.** ETFE and LDPE Film Properties.

|  | ETFE | LDPE |
| --- | --- | --- |
| Relative Permittivity | 2.6 | 2.3 |
| Electrical Conductivity | $10^{-4}$ S/m | $10^{-4}$ S/m |
| Young's Modulus | $0.66 \cdot 10^9$ Pa | $10^9$ Pa |
| Poisson's Ratio | 0.42 | 0.46 |
| Density | 1700 kg/m<sup>3</sup> | 930 kg/m<sup>3</sup> |
| Dielectric Thickness | 12.1 $\mu\text{m}$ | 12.67 $\mu\text{m}$ |

